# Meaningful patterns of information in the brain revealed through analysis of errors

**DOI:** 10.1101/673681

**Authors:** Alexandra Woolgar, Nadene Dermody, Soheil Afshar, Mark A. Williams, Anina N. Rich

**Affiliations:** Medical Research Council Cognition and Brain Sciences Unit, University of Cambridge, Cambridge, UK; Perception in Action Research Centre (PARC), Department of Cognitive Science, Macquarie University, Sydney, Australia; ARC Centre of Excellence in Cognition & its Disorders (CCD), Macquarie University, Sydney, Australia; Max Planck Institute for Human Cognitive Brain Sciences, Leipzig, Germany

## Abstract

Great excitement has surrounded our ability to decode task information from human brain activity patterns, reinforcing the dominant view of the brain as an information processor. We tested a fundamental but overlooked assumption: that such decodable information is actually used by the brain to generate cognition and behaviour. Participants performed a challenging stimulus-response task during fMRI. Our novel analyses trained a pattern classifier on data from correct trials, and used it to examine stimulus and rule coding on error trials. There was a striking interaction in which frontoparietal cortex systematically represented *incorrect* rule but correct stimulus information when participants used the wrong rule, and *incorrect* stimulus but correct rule information on other types of errors. Visual cortex, by contrast, did not code correct or incorrect information on error. Thus behaviour was tightly linked to coding in frontoparietal cortex and only weakly linked to coding in visual cortex. Human behaviour may indeed result from information-like patterns of activity in the brain, but this relationship is stronger in some brain regions than in others. Testing for information coding on error can help establish which patterns constitute behaviourally-meaningful information.

## Introduction

Successful goal-directed behaviour is thought to depend on frontoparietal cortex. In particular, certain regions of frontal and parietal cortex, comprising regions in the inferior frontal sulcus (IFS), anterior insular/ frontal operculum (AI/FO), anterior cingulate cortex / pre-supplementary motor area (ACC/pre-SMA) and intraparietal sulcus (IPS), are recruited for a wide variety of tasks (e.g. ^1,2,3^). These ‘multiple-demand’ (MD) regions ^4^, are widely implicated in neural models of cognitive control ^5-8^, in which they are proposed to selectively prioritise processing of task relevant information ^4^.

Much of the recent evidence for this idea comes from multivariate analysis of human neuroimaging data, in which patterns of activity are analysed to examine whether we can decode aspects of tasks or stimuli. For example, the MD regions have been shown to represent the key features of tasks such as stimuli, rules and responses (for a review see ^9^), and to adapt their representation of these features as required for successful behaviour ^10-14^. However, since these studies have focused almost exclusively on successful behaviour, it has been difficult to establish whether the decoded representations are actually meaningful for behaviour. Simply observing that patterns of activity differ between two conditions does not mean that this is ‘information’ that is available to other brain regions^15^, or ‘read out’ in behaviour (e.g. ^16,17,18^). Patterns observed during successful behaviour may be *necessary* for behaviour (i.e., without this pattern, this behaviour would not occur) or they could be *epiphenomenal* (e.g., an efferent copy of information, and not necessary for successful completion of the task). To test the critical assumption that multivariate patterns identify neural information relevant for behaviour, we used a novel extension of multivariate decoding in which we examined how patterns change on error trials.

If information coding in a particular brain region *is* necessary for behaviour, what would we predict on error trials? One possibility is that the brain region would fail to represent the necessary information. This is the logic used by Williams, et al. ^18^, who found that information about visual objects was absent from the object-sensitive lateral occipital complex (LOC) when participants made mistakes. In these data, object information was still present in early visual cortex (EVC), suggesting that multivoxel representation in EVC is not sufficient to drive behaviour. These authors concluded that behaviour was more closely linked to activity in the LOC than to that of the EVC. However, something important remains unexplained. There was no coding of object category in LOC, but the participant still generated a response, leaving it open as to how the incorrect response was generated. More direct evidence for a link between activation patterns and behaviour would be given by the *incorrect* information being encoded.

## Results

We designed a challenging stimulus-response task (**Figure 1**) that resulted in moderate error rates. On the basis of the behavioural response on each trial, we were able to classify responses as ‘correct’, incorrect due to application of the incorrect rule (‘rule error’), or incorrect resulting from another type of error (‘unspecified error’; e.g. misperception of stimulus, accidentally pressing the wrong button, guessing). Participants were correct on 78.2% of trials and made rule errors on 11.4% of trials, leaving 10.4% of trials as unspecified errors. We trained a pattern classifier using correct trials to derive the neural signature of correctly encoded rules and stimulus positions, and used this classifier to interrogate the content of the MD and visual cortex on error trials. This analysis approach now renders below-chance classification meaningful: it means that the pattern systematically reflects the *incorrect* information. For each error type, we were able to identify the correct or incorrect stimulus and rule information encoded in the activation patterns of different brain regions.

**Figure 1.**
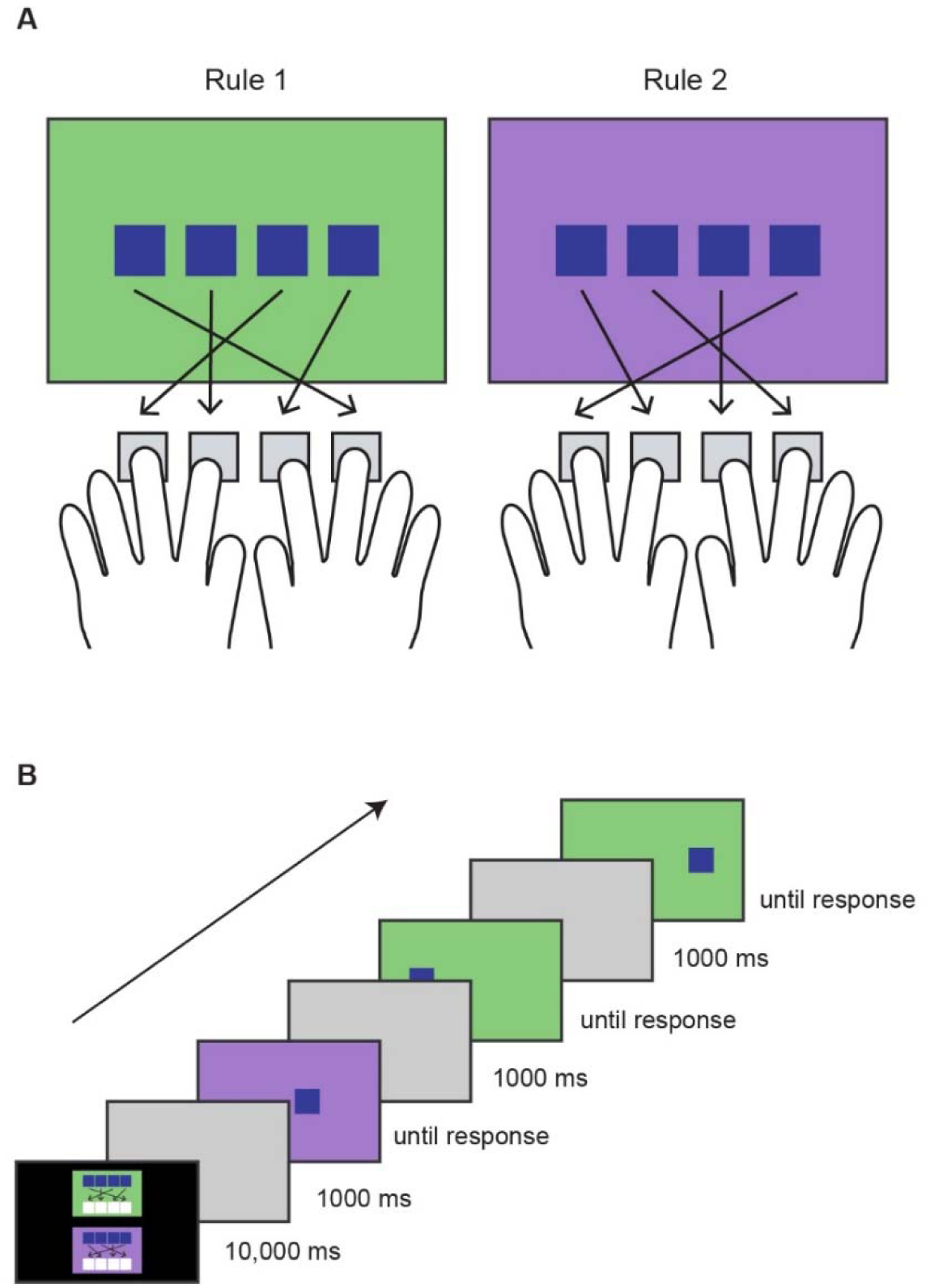
Participants observed the position of a visual stimulus on a screen and applied one of two stimulus-response mapping rules to determine which key press response should be given. (A) Each rule comprised four different position-response transformations and the two rules were mirror images of each other. Background colour indicated which rule to use on each trial (Green: rule 1; Purple: rule 2). (B) At the beginning of each block, participants were reminded of the rule mappings for 10s. Then on each trial, a single blue square was shown in one of the four possible positions against a coloured background. For the three trials shown, the correct responses would be buttons 3, 4, and 3.

There was a striking double dissociation in the MD regions (**Figure 2**). When participants made rule errors, the patterns of activation consistently reflected the *incorrect rule* (classification accuracy = 45.8%, 95% confidence intervals [41.8, 49.9], t(_21_) = −2.12, p = .046 two-tailed, Cohen’s d = −0.45; classification accuracy below 50% indicates coding of the incorrect rule). On these trials, stimulus information was encoded correctly (56.2%, [52.1, 60.2], t(_20_) = 3.15, p = .005 two-tailed, d = 0.69). For unspecified errors, we saw the reverse: now frontoparietal cortex coded the correct rule (56.1%, [50.1, 62.1], t(_16_) = 2.14, p = .048 two-tailed, d = 0.52), but incorrect stimulus (44.7%, [39.9, 49.4], t(_18_) = −2.38, p = .028 two-tailed, d = −0.55).

**Figure 2.**
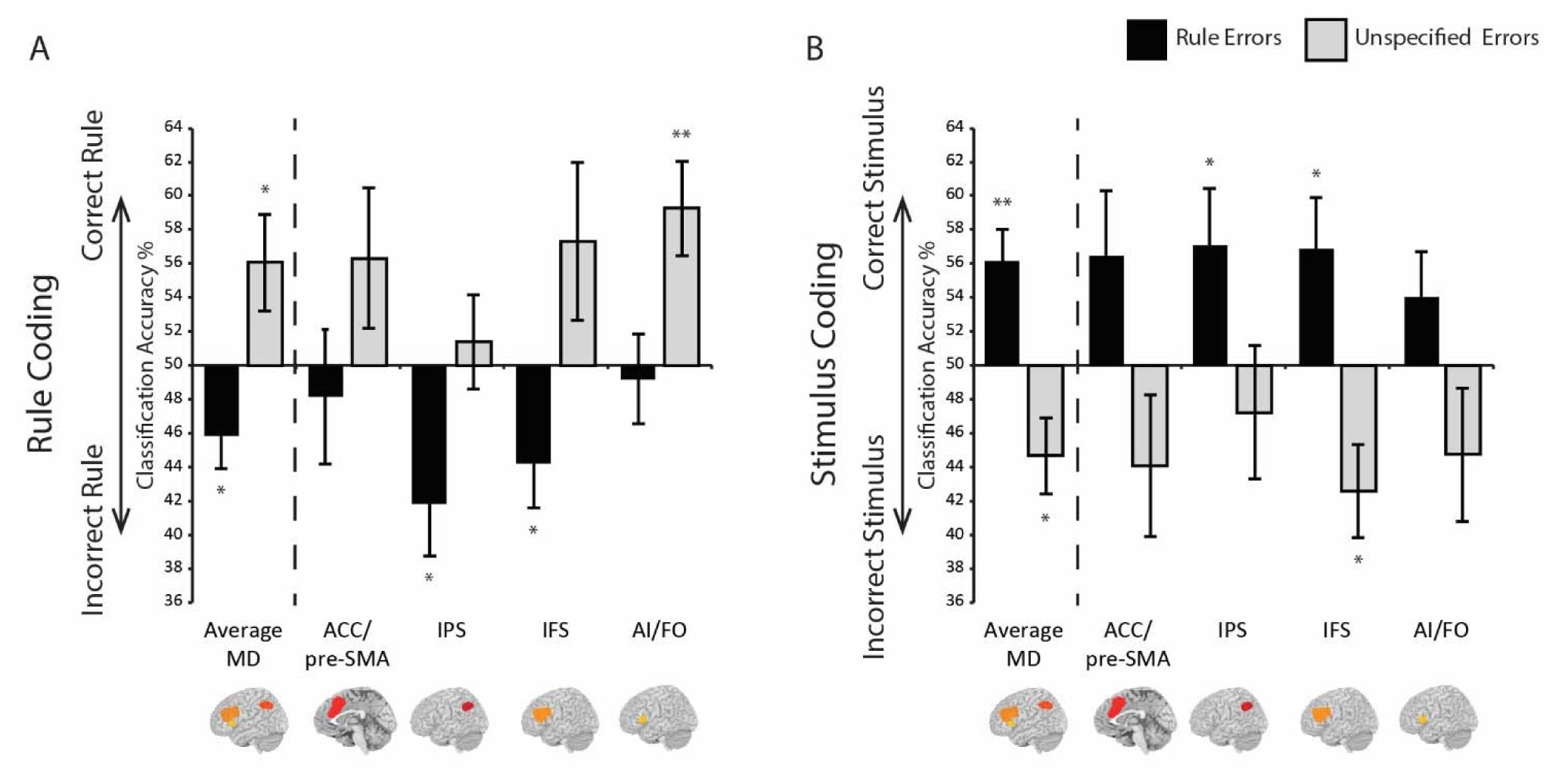
Multivoxel coding in the MD network of (A) rule and (B) stimulus information on rule error (dark bars) and unspecified error (light bars) trials. A linear support vector machine was trained on data from correct trials to establish the multivoxel pattern coding for each rule and stimulus. This classifier was then tested on data from rule error and unspecified error trials. Note that below chance (50%) classification is interpretable: below chance classification of rule indicates coding of the alternate (incorrect) rule, below chance classification of stimulus indicates coding of the alternate (incorrect) stimulus. Error bars indicate standard error. Asterisks indicate classification significantly different from chance in a two-tailed t-test against chance (50%) * p < 0.05, ** p < 0.01

We further tested whether the profile of information coding was significantly different for the two types of errors by entering the classifier accuracy scores into a repeated measured ANOVA with factors *Feature* (Rule coding, Stimulus coding), *Error Type* (Rule error, Unspecified error) and *Region* (ACC/pre-SMA, IPS, IFS, AI/FO, data collapsed over hemisphere). This showed a significant interaction between *Feature* and *Error type* (F(1,15) = 12.69, p = .003, partial eta-squared ηp^2^ =.458), indicating a different pattern of rule and stimulus information coding between rule and unspecified errors. There was no three-way interaction (F(3,45) = 0.355, p = .786, ηp^2^=.023). These results demonstrate a close link between MD coding and behaviour. On rule errors, MD activity encoded the *correct stimulus* but *incorrect rule*, while on unspecified errors, MD activity reflected the *correct rule* but *incorrect stimulus.*

A different pattern of results was apparent in visual cortex (**Figure 3**). In EVC and LOC, stimulus information was correctly encoded on rule errors (EVC: 62.3% [53.8 70.8], t(21) = 3.02, p = 0.007 two-tailed, d = 0.64; LOC: 60.7% [54.6, 66.8], t(21) = 3.64, p = 0.002 two-tailed, d = 0.78), but was absent on unspecified errors (EVC: 50.8% [42.8, 58.8], t(18) = 0.21, p = 0.84 two-tailed, d = 0.05; LOC: 51.2% [45.3 57.2], t(18) = 0.44, p = 0.67 two-tailed, d = 0.10). Thus in contrast to the MD system, the visual cortex did not encode the *incorrect* stimulus on unspecified errors, instead we failed to find any coding of stimulus information here. Rule coding was not significantly different from chance in the visual cortex for either type of error (all *p*s > 0.30, two-tailed).

**Figure 3.**
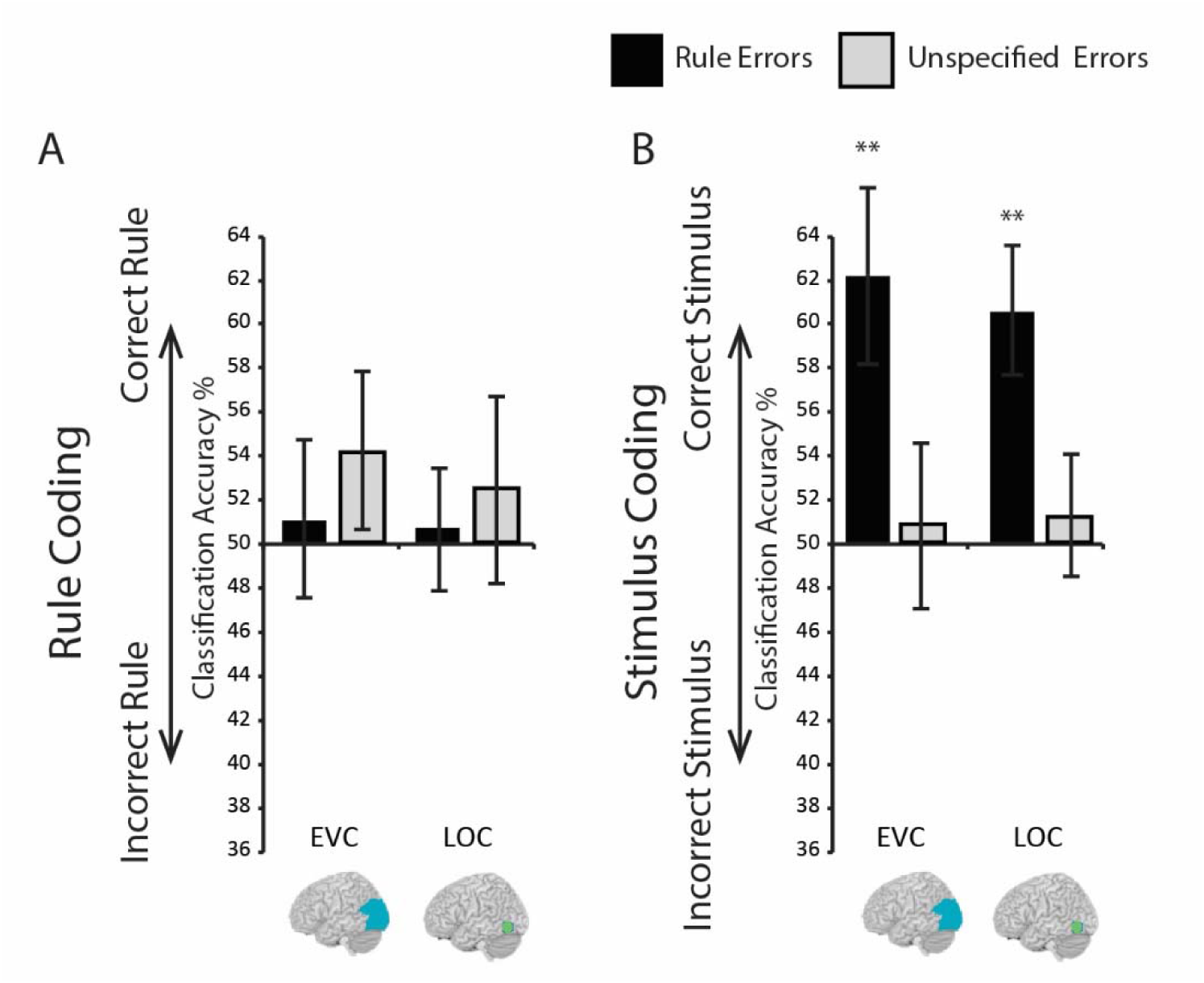
Multivoxel coding in visual ROIs of (A) rule and (B) stimulus information on rule error (dark bars) and unspecified error (light bars) trials. Conventions as in Figure 2. Stimulus position was encoded correctly when participants made rule errors, but could not be discriminated when they made unspecified errors. Unlike the MD system (Figure 2), there was no evidence for coding of the incorrect stimulus in the visual ROIs.

## Discussion

The results reveal a clear relationship between what information is coded in the MD regions and the type of behavioural error a participant makes. Behaviour appears to be more closely linked to information coding here than visual areas, consistent with a large literature suggesting a key role for these regions in cognitive control. We know, for example, that these brain regions are active for a wide range of cognitive demand (e.g. ^1,2^), and that we can decode a range of information from their activity patterns^9^. One influential theory proposes that these regions adjust their function to code information as needed for current behaviour^4^, and indeed, multivoxel codes in these regions adjust in response to a variety of task-manipulations^10-13^. Moreover, neuropsychology has long suggested a causal role for prefrontal regions in supporting meaningful goal-directed behaviour (e.g. ^19^), with recent work showing that the extent of cognitive deficit is linearly predicted by the extent of damage to the MD network^20,21^. Previously, however, it has been difficult to link these observations together – for example, to know whether the deficits observed after brain lesions are related to changes in multivoxel patterns. Here we provide correlative evidence linking multivoxel patterns in these regions to behaviour. Although we still cannot infer causality, our approach examines which multivariate patterns are predictive of behaviour, and the data underscore the importance of the MD regions in determining behavioural outcome.

We did not observe coding of the incorrect stimulus in the visual cortex on error trials: on ‘unspecified’ errors, there was no decodable information discriminating the stimuli. Therefore, the visual cortex pattern *integrity* predicted success and failure in behaviour, but, on error, we found no evidence that visual cortex coded the information used to generate the behavioural response. In contrast, for each type of error, MD patterns represented the incorrect stimulus or rule information corresponding to the incorrect behavioural response. Lacking the same evidence from EVC or LOC, we suggest the link between information coding and behavioural response here is less direct than in the MD system.

A previous study^18^ also reported that pattern integrity in LOC predicted behavioural success on a difficult object discrimination task. These authors found that stimulus information was absent on error and concluded that LOC patterns are ‘read out’ in task performance. However, this can only be concluded if LOC patterns show encoding of incorrect information, rather than reducing to noise. In our study, by training on correct trials to establish the neural signature of correctly coded information, and using this to decode error trials, we can be more specific about *what* is coded on error. Using this measure, we found that both EVC and LOC patterns resembled correct information on rule errors, but did not consistently resemble either the correct or incorrect information on unspecified errors. As with any null effect, we must be cautious. The analysis could be underpowered, although we could detect coding of the same information in the same regions on rule error trials which were very similarly powered. fMRI will also miss activity that is not extended in time, and chance decoding could reflect an average of some stimulus driven (correct) responses mixed with top-down (incorrect) influences. So we cannot conclude that the information is absent on error, but only that it does not manifest in the same way as on correct trials. Our data therefore suggest a more nuanced view: failure to represent stimulus information (in the same way as on correct trials) in the visual cortex appears to predict that the person will make an error, but at the same time, the specific behaviour that occurs cannot be directly linked to (or ‘read out’ from) the decodable information in either EVC or LOC.

The neural data point to a failure of stimulus perception on ‘unspecified’ error trials. The design of our study meant we could not independently verify when our unspecified errors followed from an incorrect perception of the stimulus rather than guesses or accidental response errors (e.g., knowing the answer but pressing the wrong button). Despite this, the representation of the incorrect stimulus in the MD system in the absence of stimulus information in the visual system suggests that the MD representation of the incorrect stimulus may have been internally generated. There are similar results when stimuli are ambiguous^22^, invisible (e.g. ^23^) or in the context of self-initiated movements or free decisions (e.g. ^24,25^), but this is the first time we see it for unambiguous and easily visible stimuli. Of course, the MD response could reflect input from weak activity in visual cortex that our methods cannot decode, or subtle contextual information such as choice history^26^; we cannot distinguish these possibilities here. Nonetheless, the dissociation between MD and visual cortex is interesting for two reasons. First, patterns reflecting visual stimuli tend to show *stronger* decoding in visual cortices than in frontoparietal cortices, making ours an unusual pattern. Second, it indicates that any feedback or re-entrant processing between the MD and visual systems (e.g. ^27,28^) was insufficient to drive decodable patterns in EVC or LOC (c.f. ^29,30,31^). Thus, decodable representation in visual cortex is not necessary for coding in MD regions or for a behavioural response to be generated. Although feedback from the MD regions may bias or support representation in the visual cortices (e.g. ^28,32^), behaviour appears to depend on representation in the MD regions themselves.

This work validates a novel variation of MVPA to examine the information on error trials: by training a classifier on plentiful correct trials, we can interrogate representational content on the less frequent incorrect trials. This approach has great potential for any question where the key condition has fewer trials than that usually required for MVPA (e.g., what aspects of processing are impaired after brain injury; what ‘lapses’ during vigilance; where errors occur within processing streams across multiple brain networks). A particularly exciting research direction may be to combine disruptive brain stimulation with these decoding techniques, to understand the causal relations between a stimulated brain area, multivoxel coding, and behaviour.

We show that activity patterns in the MD regions predict the error an individual will make. Rather than simply reducing to noise when participants make mistakes, as in visual cortex, frontoparietal activity patterns show systematic coding of incorrect information, which is in turn diagnostic of the particular error. This demonstrates the critical role of these regions in determining success or failure and draws a clear link between information coding in frontoparietal cortex and behaviour. More broadly, these results show the importance of testing for systematic links between decodable information and behaviour, in order to establish which “information” is meaningful.

## Online Methods

### Participants

Twenty-two right-handed participants (14 female, 8 male, mean age 24.9 years, SD 4.51), with normal or corrected-to-normal colour vision, took part in this study. All participants gave written informed consent and were reimbursed for their time. The study was approved by the Macquarie University Human Research Ethics Committee.

### Task

Participants performed a challenging stimulus-response task that we have used previously to separate coding of stimulus, rule and response information ^10,11^. On each trial participants identified the horizontal position of a blue square and applied one of two stimulus-response transformation rules (rule to use cued by background colour) to generate a button-press response. For further details of stimuli, task and procedure, see Woolgar, et al. ^10^. Participants learnt and performed four rules, but only the data from the two more challenging rules (shown in **Figure 1**), on which participants made a substantial number of mistakes, are analysed here. The analysis of correct trials has been published elsewhere ^10^.

### Acquisition

As described in Woolgar, et al. ^10^, we acquired fMRI scans using a Siemens 3 Tesla Verio scanner with 32-channel head coil, at the Macquarie Medical Imaging facility in Macquarie University Hospital, Sydney, Australia. We used a sequential descending T2*-weighted echo planar imaging (EPI) acquisition sequence with the following parameters: repetition time (TR), 2000 ms; echo time (TE), 30 ms; 34 oblique axial slices of 3.0 mm slice thickness with a 0.7 mm interslice gap; in-plane resolution, 3.0 × 3.0 mm; field of view, 210 mm; flip angle 78 degrees. We also acquired T1-weighted MPRAGE structural images for all participants (resolution 1.0 × 1.0 × 1.0 mm).

We presented stimuli using Matlab with Psychophysics Toolbox-3 ^33,34^, back-projected onto a screen viewed through a head-coil mounted mirror in the scanner. Participants performed 2 min blocks of the rules described here alternating with blocks of two ‘easy’ rules described in Woolgar, et al. ^10^ and not analysed here. Participants performed two EPI acquisition runs each consisting of eight blocks of trials and lasting 19 min 12 sec. Block order was counterbalanced within participants across runs, and run order was counterbalanced between participants.

At the start of each block, a graphical depiction of the two rules was displayed for 10 sec, after which the screen was grey for 1000 ms. Within each block, the eight stimuli (four positions * two background colors) were presented in random order. Stimuli remained visible for 4000 ms or until the participant responded. There was an inter-trial-interval of 1000 ms between response and display of the next stimulus, during which time the screen was grey (**Figure 1**). Block length was fixed at 2 min, in which time participants completed a varying number of trials (mean ± SD total number of trials over the 8 blocks analyzed here was 294.36 ± 95.05). At the end of each block, participants were shown a blank screen (1000 ms), the message “End of Block” (1000 ms), a blank screen (500 ms), feedback (% correct and average reaction time) for 4000 ms, and then a further blank screen (5000 ms).

### Preprocessing

Preprocessing was carried out as described in Woolgar, et al. ^10^. EPI images were spatially realigned to the first image, slice-time-corrected with the first slice as the reference and smoothed with a 4mm FWHM Gaussian kernal. The timecourse of each voxel was high-pass filtered with a cut off of 128 sec. The structural image was co-registered to the mean EPI image and normalised so as to derive normalisation parameters for deforming template space ROIs to native space.

### ROIs

We used the MD ROIs from our previous work ^11,12,21,35,36^. The seven MD ROIs were left and right IFS (centre of mass in MNI152 space: +/−38, 26, 24; volume: 17 cm^3^), left and right AI/FO (+/−35, 19, 3; 3 cm^3^), left and right IPS (+/−35, −58, 41; 7 cm^3^), and bilateral ACC/pre-SMA (0, 23, 39; 21 cm^3^). Left and right EVC ROIs (−13, −81, 3; 16, −79, 3; 54 cm^3^), were defined as Brodmann’s Areas 17 and 18, from the template available with MRIcro ^37^. Left and right LOC (+/−42 −70 −9, 4 cm^3^) was a spherical (radius: 10mm) ROI centred on co-ordinates showing maximal responses to objects over scrambled shapes and textures in the literature ^38^. We deformed these template space ROIs to the native space of each participant using the parameters derived from normalising the structural.

### First level model

We used a General Linear Model to estimate the BOLD response associated with different task events (stimulus positions, rules and responses) for correct, rule error and unspecified error trials, in each block separately. To account for trial-by-trial variation in time-on-task ^39-41^, each event was modelled using a box car function lasting from stimulus presentation until response, or the 4000ms for which the stimulus was visible if participants failed to respond. Each trial contributed to the estimation of 3 regressors (one stimulus, one rule and one response) and regressors were convolved with the haemodynamic response function of SPM (Wellcome Department of Imaging Neuroscience, London, UK; www.fil.ion.ucl.ac.uk).

### Decoding representational content on error trials

We used multivariate pattern analysis (MVPA) to examine the multivoxel representation of stimulus positions and stimulus-response mapping rules on ‘rule error’ and ‘unspecified error’ trials separately. For the analysis of stimulus positions, we compared inner with outer positions (which have equal contribution from the two rules and the four button press responses) as in our previous work ^10,11,36^. MVPA was implemented using The Decoding Toolbox ^42^ version 3.99.

For each participant and each ROI, we first trained a linear pattern classifier (LibSVMC, cost parameter C = 1) using data from the correct trials, to establish the neural signature of correctly encoded stimulus, rule and response information. For example, for the analysis of rule coding, we extracted the pattern of activation (beta estimate) associated with each rule on correct trials, in each of the 8 task blocks separately (16 betas: 2 rules * 8 blocks). This formed the dataset on which the pattern classifier was trained to distinguish between the two rules. Next, we tested the extent to which the decision boundary from this classifier cross-generalised to the unseen data from error trials. First, we tested the classification of the two rules on rule error trials. For this, we extracted the multivoxel vectors corresponding to each of the two rules that were presented on rule error trials in each of the blocks. In some cases, there were more multivoxel vectors for one rule than for the other (for example, if a participant did not happen to make a rule error in rule 1 in one of the blocks, they would not have a beta estimate for rule 1 in that block and would therefore have one fewer vectors for rule 1 than rule 2). When this happened, we ensured that the test set was balanced (equal numbers of rule error estimates for rule 1 and 2), by excluding the corresponding block from the rule with more estimates. We additionally excluded participants for whom the number of blocks per condition in the test set was less than 2. For the analysis of rule in rule error, no participants needed to be excluded and the mean (over participants) number of testing blocks retained was 5.86 (SD = 1.67) blocks for each rule. Next, we repeated the same decoding analysis for unspecified error trials (2 rules * up to 8 blocks; 5 participants excluded, mean number of testing blocks retained was 5.88 (SD = 1.83) for each rule). Note that this decoding scheme means that below chance classification accuracies are interpretable. Above chance classification signifies that the rules on error trials were correctly classified according to the neural signature derived from the correct trials. Below chance classification signifies that rules were consistently misclassified: trials where rule 1 were presented were classified as belonging to the class of activation patterns representing rule 2 and *vice versa*. Thus, below chance classification signifies consistent coding of the incorrect rule. Classification close to chance (50%) signifies no consistent information to distinguish between the two rules. Finally, the entire procedure was repeated to examine the coding of position information. For this, the following numbers of blocks were retained in the analysis: rule error: one participant excluded, mean number of testing blocks = 5.24 (SD = 1.70); unspecified error: 3 participants excluded, mean number of training blocks = 5.15 (SD = 1.70).

To assess whether any stimulus and rule information was encoded on error trials, we compared classification accuracies to chance (50%) on rule and unspecified errors separately, using two-tailed t-tests. Our unusual analysis scheme means that classification accuracies are not bounded at 50% (both above and below chance is interpretable), making parametric tests suitable^43^. We report mean classification with 95% confidence intervals, and effect size, Cohen’s d, calculated as the mean difference from chance (50%) divided by the standard deviation of the group mean classification accuracy. Additionally, to test for statistical differences in coding between the two types of error, classification accuracies for each person were entered into an ANOVA with factors *Feature* (stimulus position, rule), *Error Type* (rule error, unspecified error), and *Region* (ACC/pre-SMA, IPS, IFS, AI/FO, data collapsed over hemisphere). We report the *F* statistic and corresponding two-tail *p* value for two-way (*Feature*\**Error Type*) and three-way (*Feature*\**Error Type*\**Region*) interactions of interest, as well as effect sizes using partial eta-squared.

## Acknowledgements

This work was funded under the Australian Research Council (ARC)’s Discovery Projects funding scheme (DP12102835 and DP170101840). AW was supported by an ARC Fellowship (FT170100105) and MRC (U.K) intramural funding SUAG/035/RG91365.

## Author Contributions

A.W. designed the study, acquired the data, devised and carried out the analyses and wrote the original draft of the paper. N.D. assisted with analysis. S.A. assisted with data acquisition. M.A.W. contributed to the study design and interpretation. A.N.R. supervised the project and contributed to study design, interpretation of results, and write-up.

## References

1. Duncan, J. & Owen, A. M. Trends Neurosci 23, 475–483, (2000).

2. Fedorenko, E., Duncan, J. & Kanwisher, N. Proc Natl Acad Sci 110, 16616–16621, (2013).

3. Dosenbach, N. U. et al. Neuron 50, 799–812, (2006).

4. Duncan, J. Trends Cogn Sci 14, 172–179, (2010).

5. Dehaene, S. & Naccache, L. Cognition 79, 1–37, (2001).

6. Miller, E. K. & Cohen, J. D. Annu Rev Neurosci 24, 167–202, (2001).

7. Corbetta, M. & Shulman, G. L. Nature Reviews Neuroscience 3, 201–215, (2002).

8. Cole, M. W. & Schneider, W. Neuroimage 37, 343–360, (2007).

9. Woolgar, A., Jackson, J. & Duncan, J. J Cogn Neurosci 28, 1433–1454, (2016).

10. Woolgar, A., Afshar, S., Williams, M. A. & Rich, A. N. J Cogn Neurosci 27, 1895–1911, (2015).

11. Woolgar, A., Hampshire, A., Thompson, R. & Duncan, J. J Neurosci 31, 14592–14599, (2011).

12. Woolgar, A., Williams, M. A. & Rich, A. N. Neuroimage 109, 429–437, (2015).

13. Jackson, J., Rich, A. N., Williams, M. A. & Woolgar, A. J Cogn Neurosci 29, 310–321, (2017).

14. Li, S., Ostwald, D., Giese, M. & Kourtzi, Z. J Neurosci 27, 12321–12330, (2007).

15. de-Wit, L., Alexander, D., Ekroll, V. & Wagemans, J. Psychonomic bulletin & review, 1–14, (2016).

16. Carlson, T. A., Ritchie, J. B., Kriegeskorte, N., Durvasula, S. & Ma, J. J Cogn Neurosci 26, 132–142, (2014).

17. Ritchie, J. B. & Carlson, T. A. Front Neurosci 10, 190, (2016).

18. Williams, M. A., Dang, S. & Kanwisher, N. G. Nat Neurosci 10, 685–686, (2007).

19. Luria, A. R., (Tavistock, 1966).

20. Woolgar, A., Duncan, J., Manes, F. & Fedorenko, E. Nature Human Behaviour, (2018).

21. Woolgar, A. et al. Proc Natl Acad Sci 107, 14899–14902, (2010).

22. Roy, J. E., Buschman, T. J. & Miller, E. K. J Cogn Neurosci 26, 1283–1291, (2014).

23. Bode, S., Bogler, C., Soon, C. S. & Haynes, J. D. Neuroimage 59, 1924–1931, (2012).

24. Jenkins, I. H., Jahanshahi, M., Jueptner, M., Passingham, R. E. & Brooks, D. J. Brain : a journal of neurology 123, 1216–1228, (2000).

25. Pesaran, B., Nelson, M. J. & Andersen, R. A. Nature 453, 406, (2008).

26. Bode, S. et al. Neuroscience & Biobehavioral Reviews, (2014).

27. Di Lollo, V., Enns, J. T. & Rensink, R. A. Journal of Experimental Psychology: General 129, 481, (2000).

28. Desimone, R. & Duncan, J. Annu Rev Neurosci 18, 193–222, (1995).

29. Stokes, M., Thompson, R., Cusack, R. & Duncan, J. J Neurosci 29, 1565–1572, (2009).

30. Reddy, L., Tsuchiya, N. & Serre, T. Neuroimage 50, 818–825, (2010).

31. Cichy, R. M., Heinzle, J. & Haynes, J. D. Cereb Cortex 22, 372–380, (2012).

32. Duncan, J. & Miller, E. K. in Principals of frontal lobe function (eds D. T. Stuss & R. T. Knight) 278-291 (Oxford University Press, 2002).

33. Brainard, D. H. Spat Vis 10, 433–436, (1997).

34. Kleiner, M., Brainard, D. & Pelli, D. Perception 36, (2007).

35. Woolgar, A., Bor, D. & Duncan, J. J Cogn Neurosci 25, 1542–1552, (2013).

36. Woolgar, A., Thompson, R., Bor, D. & Duncan, J. Neuroimage 56, 744–752, (2011).

37. Rorden, C. & Brett, M. Behav Neurol 12, 191–200, (2000).

38. Thompson, R. & Duncan, J. NeuroImage 48, 436–448, (2009).

39. Woolgar, A., Golland, P. & Bode, S. Neuroimage 98, 506–512, (2014).

40. Todd, M. T., Nystrom, L. E. & Cohen, J. D. NeuroImage 77, 157–165, (2013).

41. Grinband, J., Wager, T. D., Lindquist, M., Ferrera, V. P. & Hirsch, J. NeuroImage 43, 509–520, (2008).

## Methods only references

42. Hebart, M. N., Görgen, K. & Haynes, J.-D. Frontiers in Neuroinformatics 8, 88, (2015).

43. Allefeld, C., Görgen, K. & Haynes, J.-D. Neuroimage 141, 378–392, (2016).

